# Airspace miR-146a levels in ventilated patients decrease with age and correlate with mortality

**DOI:** 10.64898/2026.06.03.728752

**Authors:** Ian D Bentley, Anusha Kapoor, Nondyce Gulick, Maya Langenecker, Laura A Leuenberger, Eric D Morrell, Joesph S Bednash, Carmen Mikacenic, Ciara M Shaver, Joshua A Englert

## Abstract

The acute respiratory distress syndrome is a heterogenous syndrome characterized by the rapid development of respiratory failure. Nearly 40% of patients who develop ARDS will die, and there is growing interest in identification of biomarkers to identify patients at risk of death and/or inform treatment decisions. Most prior work on biomarkers in ARDS has focused on the plasma compartment, but there is concern that circulating biomarkers may not reflect alveolar pathobiology. The anti-inflammatory microRNA-146a has been shown to be upregulated in inflammatory cells in human bronchoalveolar lavage fluid, but it is not known if these levels correspond with outcomes. We measured miR-146a expression by digital droplet PCR in human biospecimens from four different cohorts of patients with respiratory failure requiring mechanical ventilation – two plasma cohorts, one bronchoalveolar lavage cohort, and one heat moisture exchange (HME) filter fluid cohort. We found that miR-146a was detectible in plasma, bronchoalveolar lavage fluid, and HME fluid. However, only when measured in the alveolar space, was miR-146a expression significantly lower in older adults and those who died. It did not correlate with outcomes when measured in plasma. To our knowledge, this is the first report that nucleotides can be measured in HME fluid and builds upon expanding literature that circulating biomarkers may not reflect complex biology of the alveolar microenvironment during ARDS.

## To The Editor

The acute respiratory distress syndrome (ARDS) is a heterogenous syndrome characterized by the rapid development of non-cardiogenic pulmonary edema with hypoxemia.^1^ Approximately 10% of patients admitted to the Intensive Care Unit (ICU), and a quarter of intubated patients, develop ARDS.^2^ Nearly 40% of patients who develop ARDS will die, and increasing age is an independent risk factor for death.^2,3^ In the last decade, a number of circulating biomarkers have been identified to classify ARDS patients into either “hypoinflammatory” or “hyperinflammatory” phenotypes, and this has been shown to inform response to treatment.^4^ However, there is conflicting evidence on the degree to which circulating biomarkers reflect biology in the alveolar microenvironment.

Micro-RNAs (miR) are small non-coding RNA species that regulate gene expression. Our lab has demonstrated that microRNA-146a (miR-146a), an anti-inflammatory miR, is upregulated in inflammatory cells from human bronchoalveolar lavage (BAL) during mechanical ventilation but is not known if levels of miR-146a in airspace fluid correlate with clinical outcomes.^5^ Traditionally, biomarkers in the alveolar space, including miRs, have been measured in BAL, but this requires bronchoscopy - an invasive procedure not tolerated by all patients. The collection of fluid from exhaled breath via heat moisture exchange (HME) filters is a non-invasive method of sampling the distal airspace,^6^ but it is unknown if nucleic acid biomarkers are detectable in HME fluid. Based on our preclinical data showing that miR-146a mitigates lung injury, we hypothesized that airspace levels of miR-146a would correlate with clinical outcomes in patients with respiratory failure and that miR-146a can be detected in HME fluid.

We analyzed human biospecimens from four cohorts of patients with respiratory failure requiring mechanical ventilation: (1) plasma from 86 patients enrolled in the Aerosolized Albuterol Versus Placebo for the Treatment of Acute Lung Injury (ALTA) trial,^7^ (2) plasma from 40 patients enrolled in the BuckICU biorepository at Ohio State,^8^ (3) BAL fluid from 147 patients who underwent clinically indicated bronchoscopy for suspicion of ventilator-associated pneumonia (VAP) at the University of Washington,^9^ and (4) HME fluid from 43 patients at Vanderbilt University Medical Center.^6^ Micro-RNAs were extracted from human biospecimens using the miRNeasy Serum/Plasma Kit (Qiagen). Complimentary DNA (cDNA) was reverse transcribed and micro-RNAs within the cDNA product were amplified in an unbiased fashion using the TaqMan Advanced miRNA cDNA Synthesis Kit (ThermoFisher). Digital droplet polymerase chain reaction (ddPCR) was performed using the QX200 (BioRad) ddPCR system and miR-146a TaqMan Advanced Assay (ThermoFisher). Data were log^10^ transformed prior to analysis (HME ddPCR data was transformed after the addition of one to correct for undetectable miR-146a in some samples) and statistical testing was performed in GraphPad Prism. Further methodologic details can be found in the supplement.

Clinical characteristics of each cohort are reported in Table 1. We found that miR-146a was detectible in plasma, however, circulating levels of miR-146a did not correlate with mortality or age in either the ALTA or BuckICU cohorts (Fig 1A-D). MiR-146a was detectible in BAL fluid from critically ill adults requiring mechanical ventilation. Lower miR-146a levels were associated with increased 28-day mortality (p=0.0185, Fig 1E). BAL miR-146a levels were also decreased in aged patients compared to younger ones (p=0.0202, Fig 1F). Furthermore, we found a negative correlation between BAL miR-146a levels and BAL IL-8 levels (Table 1). We were able to detect miR-146a levels in HME fluid, and although the trends were similar to miR-146a levels in BAL, we did not have adequate statistical power to determine whether miR-146a levels correlated with mortality or age (Fig 1G-H). To our knowledge, this is the first report that microRNAs are detectable in ventilator HME fluid, and our work builds upon previous literature that supports the use of HME fluid collection for non-invasive sampling of the distal airspace.^6^

**Table 1.** Clinical characteristics of patients in each cohort and correlation analysis between BAL Cytokine and BAL miR-146a levels.

|  | ALTA | BuckICU | UW | VUMC |
| --- | --- | --- | --- | --- |
| **Number of patients** | 86 | 40 | 147 | 43 |
| **Age, avg (yrs)** | 57 | 60 | 50 | 53 |
| **Female, n (%)** | 40 (47%) | 16 (40%) | 38 (26%) | 19 (44%) |
| **ARDS, n (%)** | 86 (100%) | 40 (100%) | 77 (52%) | 31 (72%) |
| **Cytokine** | **Pearson Coefficient (95% CI)** |  | **p-value** |  |
| IFN-γ | -0.098 (-0.2560 to 0.0659) | | 0.241 | |
| IL-1β | -0.060 (-0.2202 to 0.1036) | | 0.473 | |
| IL-6 | -0.010 (-0.1723 to 0.1526) |  | 0.904 |  |
| IL-8 | -0.213 (-0.3634 to -0.0518) |  | 0.010 |  |
| IP-10 | -0.026 (-0.1887 to 0.1371) |  | 0.752 |  |
| MCP-1 | -0.079 (-0.2393 to 0.0847) |  | 0.343 |  |

**Figure 1:**
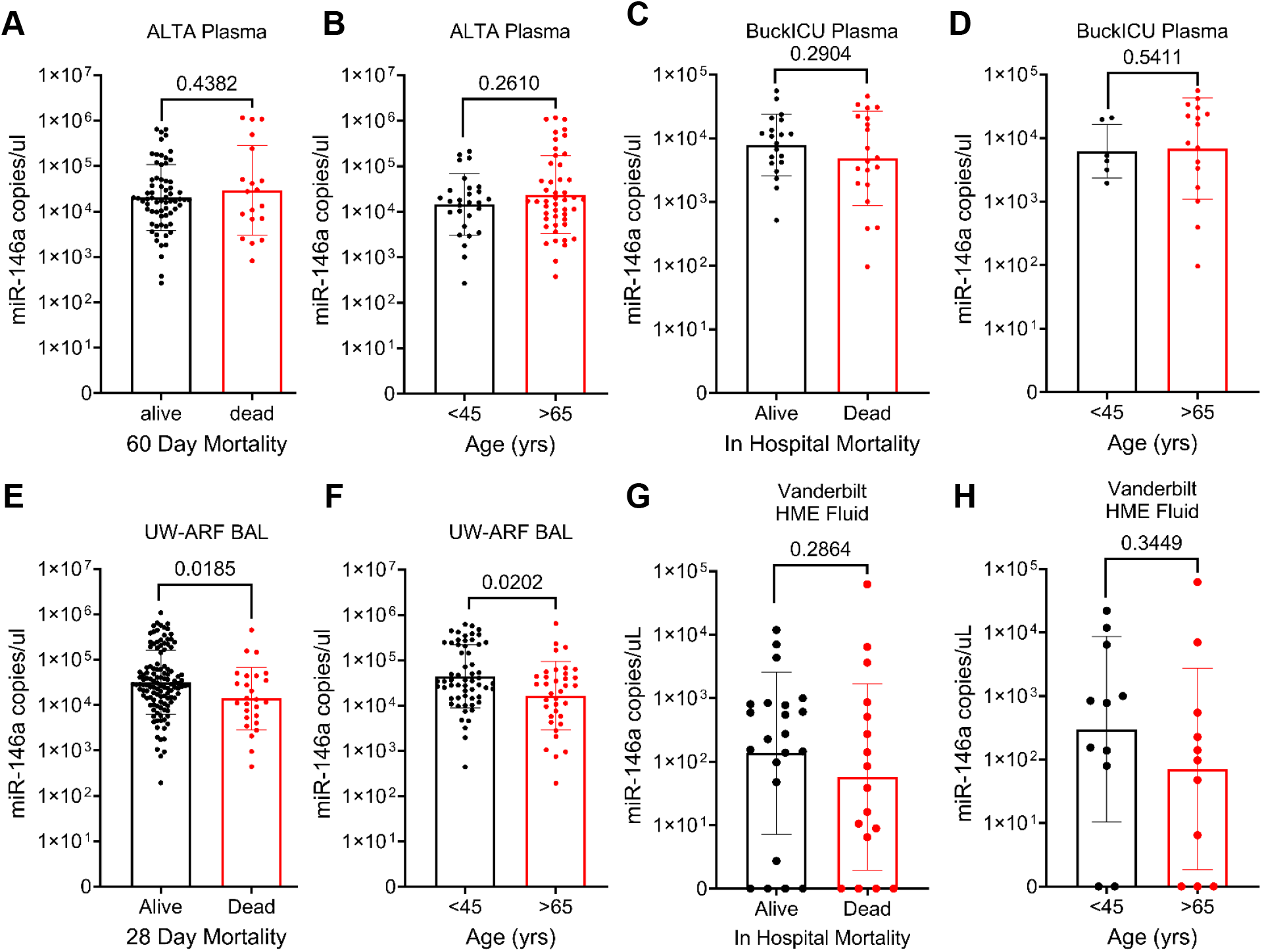
miR-146a levels in plasma, BAL and HME fluid obtained from critically ill adults requiring mechanical ventilation. All data are displayed on logarithmic scale with the geometric mean and geometric standard deviation. (A, B) miR-146a copies/uL in plasma from patients enrolled in the ARDS Network ALTA trial, correlated with mortality and age. (C, D) miR-146a copies/uL in plasma from patients enrolled in the BuckICU biorepository of biospecimens from patients with ARDS at Ohio State University, correlated with mortality and age. (E, F) miR-146a copies/uL in clinically indicated BAL performed to evaluate for suspected VAP in patients in the University of Washington medical center, correlated with mortality and age. (G, H). miR-146a copies/uL (after the addition of 1 to account for undetectable miR-146a values in 8 samples) in HME fluid from patients enrolled in the Vanderbilt HME fluid biorepository, correlated with mortality and age. All statistical testing was performed on log_10_-transformed data to account for logarithmic distribution of values. After log_10_ transformation, data was tested for normality using the Shapiro-Wilkes test and subjected to either the student’s t-test (if normally distributed) or the Mann-Whitney test (if non-normally distributed).

Previous studies have reported inconsistent correlation of airspace and plasma levels of some biomarkers (e.g. angiopoietin-2, IL6, soluble TNF receptor-1).^10^ Here, we report three clinical cohorts (2 with plasma samples and one with BAL samples) and find that only miR-146a in the airspace correlates with mortality. Interestingly, we also found that older adults, who are at increased risk of death after developing ARDS, had lower levels of miR-146a in their airspace, but not in the circulation. This finding reinforces the concept that plasma biomarker levels may not reflect the complex biology in the distal airspace of critically ill adults with respiratory failure. Interestingly, we also found an inverse correlation between BAL miR-146a and BAL IL-8 levels, consistent with the anti-inflammatory effects of miR-146a, a known negative regulator of IL-8 expression through suppression of the NF-κB/TRAF6 pathway.^5^ Our findings suggest the possibility that airspace levels of miR-146a may serve as a biomarker of inflammation in mechanically ventilated ICU patients.

Our work has several limitations. First, our airspace cohorts include a heterogenous mix of critically ill patients who have acute respiratory failure. Secondly, our biospecimens were obtained from cohorts separated in time and space, and it is possible that variations in practice or specimen processing could affect our results. However, in two disparate cohorts of plasma, one historical (ALTA) and one contemporary (BuckICU), we found that circulating miR-146a levels do not correlate with patient outcomes. These data indicate a need for future prospective studies to evaluate miR-146a (and other biomarkers) in HME fluid and plasma from patients both with and at risk for ARDS. In summary, we have shown that anti-inflammatory micro-RNAs are detectible in HME fluid from patients with respiratory failure requiring mechanical ventilation, and that these levels may correlate with clinical outcomes.

## Acknowledgements

We are grateful to the patients and their family members who participated in this research, which would not have been possible without their support. This work was prepared, in part, using Aerosolized Albuterol Versus Placebo for the Treatment of Acute Lung Injury (ALTA) research materials obtained from the National Heart, Lung, and Blood Institute’s (NHLBI) Biologic Specimen and Data Repository Information Coordinating Center and does not necessarily reflect the opinions or views of the ALTA researchers or the NHLBI. Additionally, we thank the Ohio State Comprehensive Cancer Center Genomic Core resource for their assistance with digital droplet PCR; this core resource is supported by a Cancer Center Support Grant (P30CA016058).

