## supplement for "Airspace miR-146a levels in ventilated patients decrease with age and correlate with mortality"

### SUPPLEMENTAL METHODS

Biospecimens were obtained from four cohorts of patients with respiratory failure requiring mechanical ventilation: (1) plasma from 86 patients enrolled in the Aerosolized Albuterol Versus Placebo for the Treatment of Acute Lung Injury (ALTA) trial,[1] (2) plasma from 40 patients enrolled in the BuckICU biorepository at Ohio State,[2] (3) bronchoalveolar lavage (BAL) fluid from 147 patients who underwent clinically indicated bronchoscopy for suspicion of ventilator-associated pneumonia (VAP) at the University of Washington,[3] and (4) HME fluid from 43 patients at Vanderbilt University Medical Center.[4] Total micro-RNA was extracted from equal volumes (200 uL) of human biospecimens using the miRNeasy Serum/Plasma Kit (Qiagen) per manufacturer's instructions. Complimentary DNA (cDNA) was reverse transcribed from equal volumes of total extracted micro-RNAs (2 uL per reaction) and amplified in an unbiased fashion using the TaqMan Advanced miRNA cDNA Synthesis Kit (ThermoFisher) per manufacturer's instructions to generate a miR-Amp product. The miR-Amp product was then diluted 1:10 in nuclease free $H_2O$ (Qiagen) prior to quantification by digital droplet polymerase chain reaction (ddPCR).[5]

Digital droplet PCR was performed to quantify copy numbers of miR-146a using human specific Taqman Advanced miRNA assay for miR-146a (Catalog #: A25576, Assay ID 478399_mir) and the Bio-Rad QX200 ddPCR system according to manufacturer's instructions. In brief, droplets were generated from a reaction mixture containing Taqman assay, miR-amp product and ddPCR Supermix (BioRad) and Droplet Generation Oil (BioRad) using the QX200 Droplet Generator. Each subsequent droplet randomly contains zero to one or more copies of the micro-RNA of interest based on concentration of target micro-RNA in original miR-amp product. Droplet mixture was then transferred to a 96 well plate for PCR reaction. Droplet fluorescence (or not) was determined using the QX200 Droplet Reader and final concentrations were determined using QuantaSoft Software (BioRad) by determining the fraction of negative to positively fluorescent particles.[6] Final concentrations of mir-146a in each biospecimen were determined by correction for dilution.

Clinical demographic and outcome data was available for each cohort of patients. Demographic data of interest included age, sex, race. Clinical outcome data included mortality (both in hospital and at set follow-up intervals), oxygenation (PaO2, p/f ratios, s/f ratios), ventilator settings (positive end-expiratory pressure, tidal volumes), days free of mechanical ventilation or ICU admission. In addition, BAL cytokine measurements were available for biospecimens obtained from the University of Washington. Prior to analysis, all miR-146a data was $\log_{10}$ transformed (HME data was $\log_{10}$ transformed after the addition of 1 to account for undetectable miR-146a in some samples). Graphpad prism was used to perform correlation analysis by simple linear regression for continuous variables and by the Student's t-test (if normal) or the Mann-Whitney test (if non-normal) for dichotomous variables. P-values less than 0.05 were considered significant. Multiple logistic regression was used to control for underlying patient characteristics. Data are presented on $\log_{10}$ scale.
